# Rutin derived microbiota metabolite 3,4 dihydroxybenzoic acid restores antibiotic susceptibility in XDR Gram negative pathogens with a system wide susceptibility signature

**DOI:** 10.64898/2025.12.26.696599

**Authors:** Tanim Islam, Phurt Harnvoravongchai, Alex Kidangathazhe, Sushim Gupta, Akhilesh Ramachandran, Ravi Jadeja, Joy Scaria

## Abstract

Antibiotic potentiators are a practical route to extend the utility of existing drugs against multidrug resistant Gram-negative pathogens, but most natural compound combinations remain mechanistically under defined and are rarely mapped at the level of whole cell susceptibility determinants. Here we apply a metabolite guided potentiator strategy based on gut microbiota derived breakdown products of the dietary polyphenol rutin, and we integrate potentiation phenotyping with system wide genetic susceptibility mapping to guide combination design. Three rutin metabolites, 3,4 dihydroxybenzoic acid (DHBA), 2,4,6 trihydroxybenzoic acid (THBA), and 3,4 dihydroxyphenylacetic acid (DOPAC), were screened against a CDC ARLG reference panel composed of multidrug resistant and extensively drug resistant (XDR) isolates, identifying DHBA as the most broadly active candidate, including against XDR *Klebsiella pneumoniae, Acinetobacter baumannii, Pseudomonas aeruginosa,* and *Escherichia coli*. DHBA restored susceptibility to selected antibiotic classes in resistant Gram-negative pathogens, including an approximately sixteen-fold reduction in colistin MIC in mcr 1 positive *E. coli*. To better understand the potentiation mechanism, we performed a large-scale genetic susceptibility screen of 316 *E. coli* single gene knockouts and defined a DHBA response architecture enriched for envelope and transport determinants with additional contributions from central metabolism and information processing pathways. Comparative mapping under colistin exposure revealed a distinct susceptibility architecture with limited overlap, supporting the concept that both potentiators and antibiotics engage secondary cellular systems beyond canonical primary mechanisms. In a Mini Bioreactor Array gut community model, DHBA produced a more conserved community shift than colistin. Finally, DHBA-antibiotic combinations improved outcomes in infection relevant models, including improved survival in *Galleria mellonella*, reduced intestinal burden in *Caenorhabditis elegans*, and reduced bacterial burden in an *ex vivo* porcine burn wound infection model. Collectively, these findings support a systems-based framework for developing mechanistically informed potentiator antibiotic-combinations to extend the lifespan of existing antibiotics.

## Introduction

Antimicrobial resistance (AMR) is a leading cause of mortality worldwide where 5 million deaths are associated with AMR bacterial infections(1). It is eroding the effectiveness of antibiotics used to treat bacterial infections in humans and animals(2). The greatest clinical pressure is increasingly resistant in Gram negative pathogens, where intrinsic permeability barriers and acquired resistance mechanisms converge to limit therapeutic options(3). Multidrug resistant (MDR) and extensively drug resistant (XDR) strains within the ESKAPEE pathogens *(Enterococcus spp., Staphylococcus aureus, Klebsiella pneumoniae, Acinetobacter baumannii, Pseudomonas aeruginosa, Enterobacter spp.,* and *Escherichia coli)* (4) now account for a substantial fraction of AMR associated morbidity and mortality(1, 5). Even last resort antibiotics are being compromised, exemplified by the global spread of colistin resistance(6, 7). At the same time, the pace of antibiotic discovery has not kept up with the speed of resistance evolution, leaving a narrow set of available drug classes to manage resistant infections(8). Because of these constraints, new strategies that extend the utility of existing antibiotics are needed rather than relying solely on the introduction of new agents.

Antibiotic potentiators offer a pragmatic route to restore activity of existing drugs against resistant pathogens(9). Potentiators are agents that increase antibiotic efficacy in combination, often by neutralizing resistance mechanisms or shifting bacterial susceptibility states(10). The clinical success of beta lactam antibiotics paired with beta lactamase inhibitors illustrates the value of this approach(11, 12). However, most candidate potentiators, particularly natural products, have been advanced primarily on the basis of synergy endpoints, with limited mechanistic depth(13). As a result, combination selection, dose optimization, and prediction of strain to strain variability remain largely empirical. A mechanistic framework that explains how potentiators reshape susceptibility is needed to enable rational combination design and to improve translational potential(14).

A recent shift in antibiotic research has been a move away from a strict “one drug one target” view toward cell wide susceptibility architectures(15). Antibiotic efficacy in Gram-negative bacteria is governed not only by the canonical primary target, but also by envelope access, efflux and transport, stress responses, and metabolic state(16). Increasing evidence indicates that antibiotics engage secondary cellular systems that shape tolerance, recovery, and adaptation(17, 18). For potentiators, this implies that meaningful progress requires mapping genetic determinants of susceptibility at the scale of cellular systems, rather than relying on endpoint synergy alone. Such systems level mapping can identify the modules that govern potentiation, distinguish shared versus distinct susceptibility architectures across drug classes, and guide pairing strategies that are mechanistically complementary.

Dietary polyphenols are widely encountered phytochemicals that have been explored as antimicrobial adjuvants, but their direct antibacterial effectiveness is often limited by physicochemical constraints, including size and restricted penetration across Gram-negative barriers(19, 20). In contrast, gut microbiota metabolism converts many dietary polyphenols into smaller phenolic acids(21), that occupy a more permissive chemical space for Gram-negative entry and membrane interaction(22). This metabolic conversion provides a rational discovery strategy: rather than screening parent polyphenols as broad natural products, microbiota derived metabolites can be prioritized as candidate potentiators with improved access to bacterial cellular systems. Rutin is a representative dietary polyphenol whose microbial breakdown yields several phenolic acid metabolites that are plausible potentiator candidates(23, 24).

Here, we apply a metabolite guided potentiator strategy focused on three rutin derived gut microbiota metabolites, 3,4 dihydroxybenzoic acid (DHBA), 2,4,6 trihydroxybenzoic acid (THBA), and 3,4 dihydroxyphenylacetic acid (DOPAC). We first screened these metabolites against an reference panel containing susceptible, MDR and XDR strains and identified DHBA as the most broadly active candidate, including activity against XDR isolates. We then tested whether DHBA restores susceptibility to clinically relevant antibiotics in resistant Gram-negative pathogens, establishing class selective potentiation. To define mechanism of action, we integrated membrane permeability and integrity phenotyping with a curated genetic susceptibility screen composed of large number of single gene mutants across major cellular modules in *E. coli*, and we used colistin as a comparator to distinguish convergent envelope phenotypes from divergent susceptibility architectures. As supporting translational evidence, we assessed collateral community effects in a Mini Bioreactor Array gut microbiome model and evaluated DHBA antibiotic combinations in infection relevant models, including *Galleria mellonella, Caenorhabditis elegans*, and an *ex vivo* porcine skin wound infection model. Together, this work links a microbiota derived metabolite potentiator to cell wide susceptibility mapping and provides a mechanistically informed framework for designing antibiotic potentiator combinations against resistant Gram-negative pathogens.

## Materials and methods

### Bacterial strains and reagents

All bacterial strains used in this study are listed in Supplementary Table 1. Strains were stored at −80 °C in nutrient broth supplemented with 18% (v/v) dimethyl sulfoxide. For experiments, strains were cultured in brain heart infusion (BHI) broth or Luria Bertani (LB) broth, as specified for each assay. Antibiotics used in this study included colistin (polymyxin; Sigma), kanamycin (aminoglycoside; Sigma Aldrich), cefepime (beta lactam; Sigma Aldrich), and ciprofloxacin (fluoroquinolone; Thermo Fisher Scientific). All antibiotic stock solutions, except ciprofloxacin were prepared in sterile water. Ciprofloxacin stock solution was prepared in 0.1M HCL. DHBA was dissolved in prewarmed 3% ethanol, DOPAC was dissolved in sterile water, and THBA was dissolved in dimethyl sulfoxide. All stock solutions were filter sterilized prior to use.

### MIC determinations

Minimum inhibitory concentrations (MICs) were determined by broth microdilution according to CLSI 2018 guidelines with minor modifications. Two-fold serial dilutions of each compound were prepared in BHI or LB broth and dispensed into 96 well microtiter plates (CellTreat, NY, USA). Bacterial suspensions were prepared in the corresponding broth and were added to each well-adjusted to final starting inoculum of OD 0.05. Plates were incubated at 37 °C for 18 to 24 h. MIC was defined as the lowest concentration with no visible growth. Absorbance at 630 nm was measured to quantify percent growth relative to untreated controls.

### Phenolic acid screening across a CDC reference pathogen panel

To screen phenolic acid activity against antibiotic resistant strains, we used a reference set of isolates sourced from the CDC ARLG Antibiotic Resistance Isolate Bank(25). Thirty-eight strains spanning major Gram negative and Gram positive multidrug resistant and extensively drug resistant pathogens were screened in 96 well microtiter plates. Overnight cultures grown in BHI were diluted to a starting inoculum of OD600 = 0.1 to 0.15 and treated with phenolic acids at final concentrations of 2000, 3000, and 4000 µg/mL. Four plates were prepared for each treatment and four plates were used as the common untreated control (BHI only) for DHBA and DOPAC. Four plates were used as the corresponding vehicle control (4% dimethyl sulfoxide) for THBA. Plates were incubated at 37 °C for 18 to 24 h without shaking.

Relative growth percentage (RGP) was calculated as:

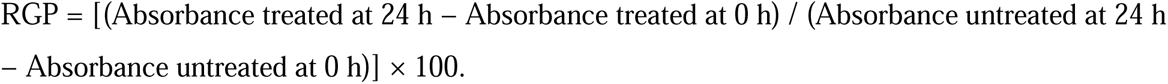

### Mini-Bioreactor Array (MBRA) system set up and treatment of fecal sample

Following approval from the South Dakota State University IRB, fecal samples were collected from healthy adult donors with no antibiotic exposure within the preceding year. Following our previously published protocol, samples were processed anaerobically within two hours of collection and stored at −80 °C until further use. A modified brain heart infusion (mBHI) medium served as the basal growth medium for all bioreactor runs(26). Mini Bioreactor Array (MBRA) units were assembled and sterilized as described previously(27–29) with minor modifications. Each reactor had a 15 mL working volume and was operated inside an anaerobic chamber (Coy Laboratory Products) maintained at 37 °C under an atmosphere of 85% N□, 10% CO□, and 5% H□. The inflow medium pH was adjusted to 6.8 and maintained constant throughout the experiment. Continuous flow was controlled using dual 24 channel Watson Marlow pumps, with inflow and outflow rates set at 1 rpm and 2 rpm, respectively, yielding an average retention time of 16 h. Magnetic stirring was maintained at 130 rpm.

Following equilibration, 300 µL of the stored fecal inoculum was introduced into each reactor. Four conditions were tested in triplicate: untreated control (mBHI only), DHBA (final concentration 3000 µg/mL), colistin (final concentration 6.25 µg/mL), and DHBA plus colistin. Treatments were administered daily from day 7 through day 14. Continuous culture was maintained for 14 days, and 1 ml aliquots were collected aseptically on day 14 for sequencing. Samples for DNA extraction and 16S rRNA sequencing were immediately frozen at −80 °C. This configuration enables stable anaerobic cultivation of complex fecal communities and supports comparative assessment of treatment associated changes in community composition and resilience over time.

### Nanopore sequencing and bioinformatic analysis of MBRA communities

DNA extraction Microbial DNA was extracted from frozen MBRA samples using the ZymoBIOMICS DNA Miniprep Kit (Zymo Research, Irvine, CA, USA) according to the manufacturer’s instructions. Samples were thawed on ice, mixed thoroughly, and processed using the kit’s mechanical lysis workflow, including bead based disruption, to support recovery of DNA from both Gram positive and Gram-negative taxa. An extraction blank was processed in parallel as a negative control. Extracted DNA was quantified using the NANODROP ONE ^©^(Thermo Scientific) and stored at −20 °C until library preparation.

### 16S rRNA gene library preparation and Nanopore sequencing

Amplicon DNA generated from 16S rRNA gene PCR was used as input for sequencing library preparation. Libraries were prepared using the Oxford Nanopore Native Barcoding Kit 96 V14 (SQK NBD114.96), which enables multiplexing of amplicon DNA, according to the manufacturer protocol. For each sample, 200 fmol of amplicon DNA was used as input for barcoding. Amplicon ends were prepared for ligation using the NEBNext Ultra II End Repair and dA Tailing Module, followed by ligation of a unique native barcode using the Native Barcoding Kit reagents. After barcode ligation, reactions were stopped with EDTA and barcoded samples were pooled, followed by bead based clean up and elution in nuclease free water. Sequencing adapters were then ligated using the kit native adapter and a quick ligation reaction, followed by bead based clean up using the Short Fragment Buffer to retain amplicon DNA. The final library was eluted in elution buffer and quantified by Qubit. The library was adjusted to 12 µL at 10 to 20 fmol for loading. Sequencing was performed on R10.4.1 flow cells (FLO MIN114) on a GridION device using MinKNOW software. The flow cell was primed using Flow Cell Flush and Flow Cell Tether, and bovine serum albumin was included in the priming mix at a final concentration of 0.2 mg/mL. Libraries were prepared for loading using sequencing buffer and library beads, then loaded through the SpotON port according to the manufacturer guidance. Base calling was performed in MinKNOW, and demultiplexing of barcoded reads was performed in MinKNOW using the SQK NBD114.96 kit setting. Demultiplexed reads were exported as FASTQ files for downstream processing.

Diversity and statistical analysis: Quality filtering and OTU generation were performed using VSEARCH (version 2.30.2). Reads were filtered using a minimum quality threshold of 0 ambiguous bases and length constraints appropriate for the expected amplicon size (400 to 1400 bp). Dereplication was performed with 98% identity and singleton handling was set to remove singletons. Chimeric sequences were removed using both de novo chimera detection (uchime de novo) and reference-based chimera detection (uchime ref) against SILVA (version 138.2). Operational taxonomic units were clustered at 97% sequence identity to generate representative sequences and an OTU abundance table. Representative sequences were classified using the VSEARCH implementation of the SINTAX algorithm with the SILVA (version 138.2) reference database. Taxonomy was summarized at the species level. Downstream statistical analyses were performed in R (version 4.3.1). Community composition plots were generated using ggplot2 (version 4.0.1). Ordination was performed using principal component analysis or principal coordinates analysis, as appropriate, implemented in FactoMineR (version 2.11) and visualized using ggplot2.

### Checkerboard interaction assays and CFU enumeration

Combinatorial interactions between DHBA and antibiotics were evaluated using two-dimensional checkerboard assays in 96 well microtiter plates. Two-fold serial dilutions of each antibiotic were prepared along the horizontal axis, and DHBA was diluted along the vertical axis. Each plate included untreated growth controls and vehicle controls matched to the DHBA solvent condition. Bacterial cultures were grown overnight in LB or BHI broth, diluted into fresh medium, and added to each well to a final OD600 of 0.05. Plates were incubated at 37 °C for 18 to 24 h. Growth inhibition was quantified by measuring absorbance at 630 nm and expressed relative to untreated controls.

To validate absorbance-based inhibition and to assess bactericidal activity in selected conditions, colony forming units (CFUs) were quantified by serial dilution and spot plating on LB agar. Wells selected for CFU enumeration included untreated controls, single agent conditions near the MIC, and combination wells corresponding to the lowest inhibitory concentrations identified by the checkerboard assay. Serial dilutions were prepared in sterile diluent and plated by spot inoculation, followed by incubation at 37 °C until colonies were countable. CFU per mL was calculated based on dilution factor and plated volume. Each assay was performed with independent biological replicates.

### Outer membrane permeability assay

Outer membrane permeability was assessed using 1-N phenylnaphthylamine (NPN; Sigma Aldrich) at a final concentration of 1 µM. *E. coli* BAA 3170 was grown to mid log phase (OD600 approximately 0.5). Cells were treated with DHBA or colistin at their respective MIC concentrations for 5 h at 37 °C with shaking at 200 rpm. Following treatment, cells were pelleted and washed twice with phosphate buffered saline. Washed cells were resuspended in phosphate buffered saline and normalized to OD600 = 0.5. NPN was added, and samples were incubated for 20 min in the dark in standard 96 well microtiter plates (Fisher Scientific). Fluorescence was measured immediately after incubation using an Infinite M200 microplate reader (Tecan) with excitation at 340 nm and emission at 485 nm. Untreated controls were included.

### Inner membrane permeability assay

Propidium iodide (PI; Thermo Fisher Scientific) was used to assess inner membrane integrity as described previously (30), with minor modifications. E. coli BAA 3170 was grown to mid log phase (OD600 approximately 0.5) and treated with DHBA or colistin at their respective MIC concentrations for 30 min at 37 °C with shaking at 200 rpm. Cells were pelleted and washed twice with phosphate buffered saline, resuspended in phosphate buffered saline, and normalized to OD600 = 0.5. PI was added at a final concentration of 10 nM, and samples were incubated for 20 min in the dark in standard 96 well microtiter plates (Fisher Scientific). Fluorescence was measured immediately after incubation using an Infinite M200 microplate reader (Tecan) with excitation at 535 nm and emission at 612 nm. Untreated and positive controls were included.

### Genetic susceptibility screening of an E. coli single gene knockout library

A panel of 316 single gene knockout strains from the Keio collection (31), was screened to identify genetic determinants of susceptibility to DHBA and colistin (Supplementary table 2). Mutants were revived in LB broth supplemented with kanamycin (25 µg/mL) and grown overnight. Cultures were then normalized to an initial OD600 of 0.5 to 0.7 and dispensed into standard 96 well microtiter plates (Fisher Scientific) at a final volume of 40 µL per well. DHBA (final concentration 1750 µg/mL) or colistin (final concentration 1.5 µg/mL) was added in LB supplemented with kanamycin. Wild type *E. coli* BW25113 and *E. coli* DH5α were included as controls. Plates were incubated at 37 °C with shaking, and absorbance at 630 nm was recorded at baseline and at multiple time points during incubation. Relative growth percentage (RGP) was calculated using the approach described above, with treated growth normalized to the corresponding untreated controls.

To characterize treatment associated growth dynamics, selected knockout strains were further evaluated using kinetic growth assays under the same media and drug exposure conditions. Overnight cultures were normalized to OD600 = 0.5, transferred to standard 96 well plates (40µL per well), and exposed to DHBA (1750 µg/mL) or colistin (1.5 µg/mL) in LB supplemented with kanamycin. Plates were incubated at 37 °C with shaking, and OD630 was recorded at regular intervals over 24 h to quantify temporal growth inhibition profiles relative to untreated controls.

### Swimming motility assay

Swimming motility was assessed using LB soft agar plates containing 0.3% agar, prepared with or without DHBA supplementation (final concentration 1750 and 2000 µg/mL). Bacterial cultures were grown to OD600 = 0.5, and 2 µL of culture was inoculated into the center of each plate by a single stab. Plates were incubated at 37 °C for 48 h under static conditions. Swimming motility was quantified by measuring the diameter of the migration zone from the point of inoculation.

### Galleria mellonella infection model

*Galleria mellonella* larvae (180 to 250 mg) were randomly assigned to four groups (n = 17 to 20 larvae per group) and infected with *E. coli* BAA 3170 (1 × 10^6 CFU per larva) by injection into the left proleg using a 10 µL Hamilton micro syringe (Hamilton 80300). At 1 h post infection, treatments were administered by injection into the right proleg using phosphate buffered saline as the vehicle, as follows: colistin (2.5 mg/kg), DHBA (500 mg/kg), or the combination of colistin and DHBA. An infected vehicle treated group served as the control. Larvae were incubated at 25 °C, and survival was monitored daily for 5 days. Death was defined as absence of movement in response to gentle stimulation and black coloration of the body.

### Pathogen clearance in Caenorhabditis elegans

Synchronized L1 stage *Caenorhabditis elegans* (N2 strain) were obtained by bleach synchronization followed by hatching in M9 buffer. The intestinal colonization assay was performed in liquid culture with 30 worms per condition. Worms were exposed to E. coli BAA 3170 in liquid media containing colistin (1 µg/mL), DHBA (2000 µg/mL), or the combination of colistin and DHBA. A no drug condition served as the control. After 2 days of exposure, worms were collected, washed to remove non-adherent bacteria, surface sterilized, and lysed. Lysates were serially diluted and plated on LB agar to quantify intestinal bacterial burdens by colony forming unit enumeration.

### Ex vivo porcine skin wound infection model

The effect of DHBA in porcine skin wounds was evaluated using an ex vivo burn wound infection model as described previously(32). Porcine skin blocks (approximately 5 cm²) were prepared and stored at −20 °C until use. Prior to infection, skin blocks were sterilized and burn wounds were generated using a preheated soldering iron fitted with an 8 mm tip (Weller, USA) applied for 20 to 40 s. Wounds were inoculated with 30 µL of *Pseudomonas aeruginosa* 2339 (10^8 CFU/mL). After a 2 h incubation at 37 °C to allow bacterial establishment, wounds were treated (100 µL per wound) with DHBA (2000 µg/mL), cefepime (4 µg/mL), or the DHBA plus cefepime combination (n = 4 wounds per condition). Following an additional 4 h incubation at 37 °C, wounds were washed twice with 50 µL phosphate buffered saline, and bacterial burden was quantified by serial dilution and plating on BHI agar plates.

## Results

### Phenolic acid screening identifies DHBA as the most active compound against MDR and XDR clinical isolates

We screened three gut microbiota derived metabolites of the dietary polyphenol rutin, DHBA, THBA, and DOPAC, selected for their lower molecular weight and potential to better satisfy bacterial entry constraints, against a CDC ARLG reference panel (Supplementary table 1) spanning multidrug resistant and extensively drug resistant clinical isolates (Fig. 1). DHBA showed the broadest inhibitory profile across the panel, with consistent suppression of growth at the tested concentrations. Notably, DHBA inhibited several isolates classified as XDR in this reference set, including XDR *K. pneumoniae* (ARLG 1129, 1139, and 1351), multiple XDR*A. baumannii* isolates (ARLG 1299, 1863, 1878, and 1913), and the VIM positive XDR *P. aeruginosa* 2339. DHBA also inhibited the XDR E. *coli* isolate 1012. In comparison, THBA produced more limited growth suppression across the panel, with the clearest inhibition appearing in a subset of Gramnegative isolates at higher concentrations, while several *Enterobacterales* remained comparatively less affected. DOPAC showed a broader concentration dependent inhibitory pattern than THBA, with more consistent suppression across many Gram-negative isolates at 3000 to 4000 µg/mL, but still weaker overall activity than DHBA at lower concentrations. Based on its broader inhibition of MDR and XDR isolates in the CDC reference panel, DHBA was prioritized for subsequent potentiation and mechanistic analyses.

**Fig. 1.**
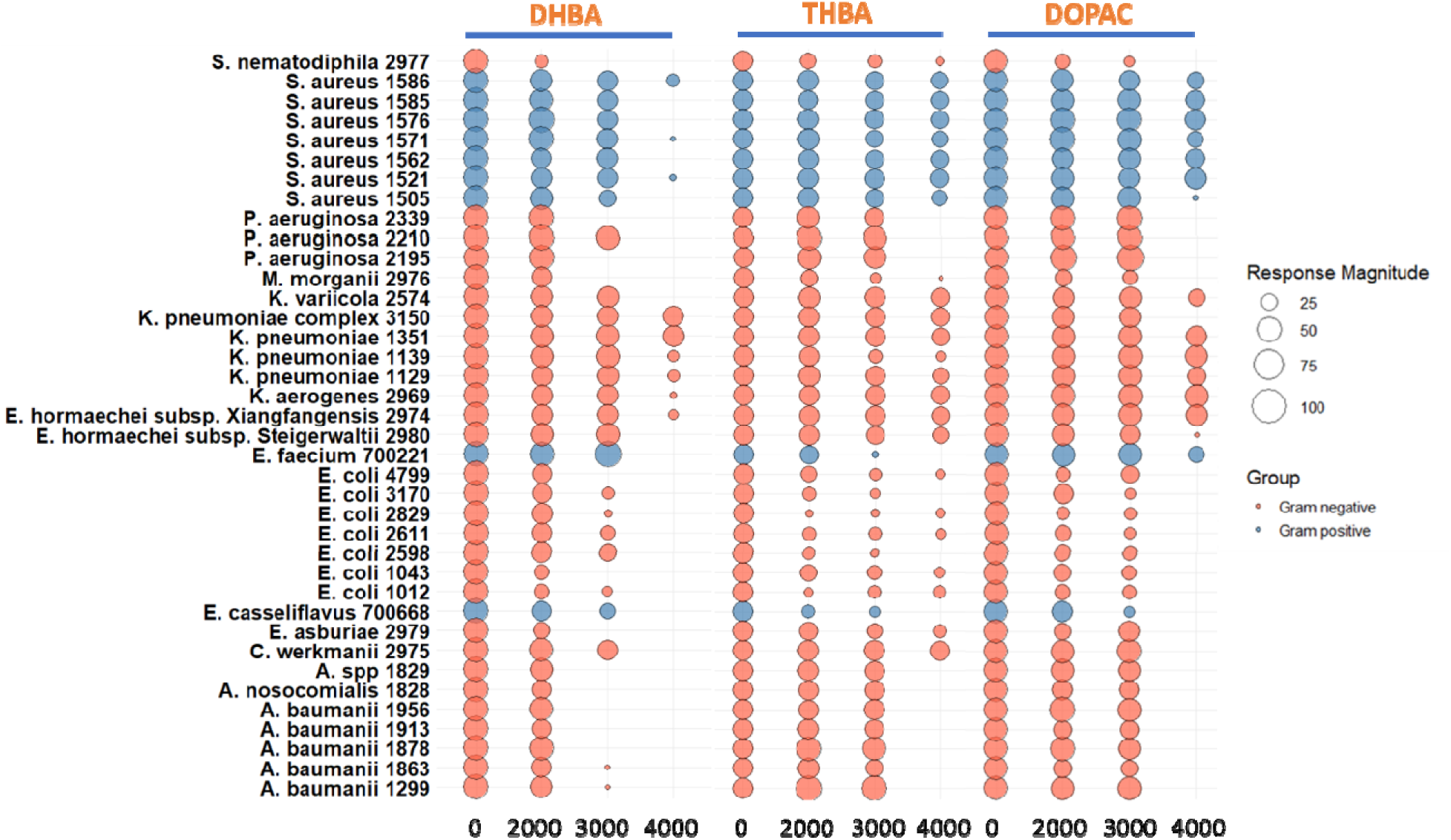
Phenolic acid screening across a CDC reference pathogen panel. Growth inhibition profiles of 38 bacterial strains representing 16 species, including Gram-positive (blue) and Gram-negative (red) pathogens, after exposure to increasing concentrations of 3,4 dihydroxybenzoic acid (DHBA), 3,4 dihydroxyphenylacetic acid (DOPAC), or 2,4,6 trihydroxybenzoic acid (THBA). Cultures were incubated for 18 to 24 h at 37 °C in BHI broth, and growth was quantified by absorbance at 630 nm. Bubble size denotes relative growth percentage relative to untreated controls (100, 75, 50, or 25%), calculated from baseline and endpoint absorbance measurements as described in Methods. Strains were sourced from the CDC ARLG Antibiotic Resistance Isolate Bank and include multidrug resistant and extensively drug resistant clinical isolates.

### DHBA restores susceptibility to multiple antibiotic classes in resistant Gram-negative pathogens

Because DHBA emerged as the most active rutin derived microbiota metabolite across the CDC reference panel, we next tested whether DHBA can function as an antibiotic potentiator and restore susceptibility in resistant Gram-negative pathogens. We evaluated DHBA in two-dimensional checkerboard assays with representative antibiotics spanning four mechanistic classes, including the last resort polymyxin colistin. For all assays, bacterial inocula were standardized to OD600 = 0.05 to enable direct comparison across conditions. DHBA showed class selective potentiation rather than a universal enhancement of all antibiotics. In *E. coli*, DHBA did not alter kanamycin activity, and the kanamycin MIC remained largely unchanged across the DHBA concentration range tested (Fig. 2a). In contrast, DHBA strongly potentiated colistin against *E. coli*, restoring susceptibility at sub inhibitory DHBA concentrations and decreasing the colistin MIC by 16-fold (Fig. 2b). DHBA also potentiated cefepime and ciprofloxacin in *E. coli*, shifting MICs into a range where growth inhibition was evident under conditions where these antibiotics alone showed limited activity at the concentrations tested (Fig. 2c–f). To test whether this potentiation extends beyond *E. coli,* we evaluated DHBA combinations in two extensively drug resistant pathogens. DHBA enhanced cefepime activity against *P. aeruginosa* 2339 and reduced the colistin MIC in *K. pneumoniae* 1129 (Fig. 2d–e). Together, these data indicate that DHBA can restore susceptibility to selected antibiotic classes in resistant Gram-negative pathogens, with potentiation patterns that are consistent with a mechanism linked to envelope associated susceptibility rather than non-specific growth inhibition.

**Fig. 2.**
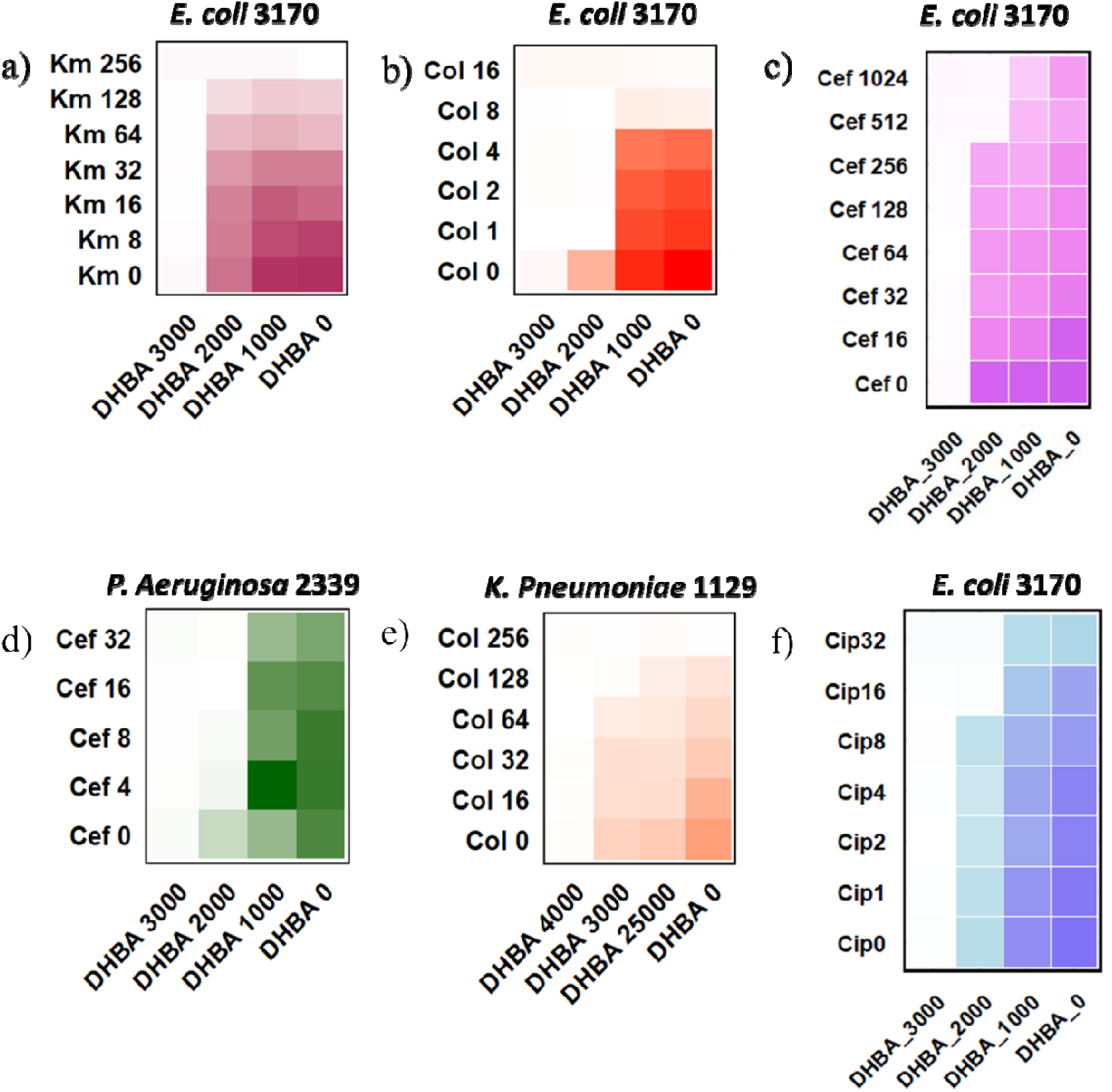
DHBA restores susceptibility to multiple antibiotic classes in resistant Gram-negative pathogens. Two dimensional checkerboard assays showing antibiotic activity across increasing concentrations of DHBA. Bacterial inocula were standardized to OD600 = 0.05, and plates were incubated for 18 to 24 h at 37 °C. Growth was quantified by absorbance at 630 nm. (a) Kanamycin activity in mcr 1 *E. coli* BAA 3170 in the presence of DHBA, showing no appreciable potentiation. (b) Colistin activity in mcr 1 *E. coli* BAA 3170, showing DHBA mediated resensitization with an approximately 16-fold reduction in colistin MIC. (c) Cefepime and (f) ciprofloxacin activity in mcr 1 *E. coli* BAA 3170, showing DHBA mediated MIC shifts. (d) Cefepime activity in XDR *P aeruginosa* 2339 in the presence of DHBA. (e) Colistin activity in XDR *K. pneumoniae* in the presence of DHBA, showing an approximately 2-fold MIC reduction. Heatmaps depict growth inhibition across the antibiotic and DHBA concentration matrix, where darker shading denotes higher growth. Values represent mean OD630 from three or more independent biological replicates.

### DHBA antibiotic combinations show bactericidal activity by CFU enumeration

To determine whether DHBA mediated potentiation corresponds to bactericidal killing rather than growth suppression alone, we quantified viable bacteria by colony forming unit (CFU) enumeration following checkerboard exposures. Colistin is a bactericidal last resort antibiotic for Gram-negative pathogens, and therefore provides a stringent benchmark for evaluating DHBA adjuvant effects. In *E. coli*, DHBA at 2000 µg/mL increased colistin associated killing in a concentration dependent manner at lower colistin exposures (1 to 4 µg/mL), resulting in a marked reduction in recoverable CFUs compared with colistin alone (Fig. 3a). At higher colistin exposure (8 µg/mL), the DHBA combination did not further decrease CFUs beyond the effect of colistin alone, consistent with a plateau in bactericidal activity under these conditions (Fig. 3a). We next tested whether DHBA can enable near complete clearance when paired with additional antibiotic classes. In *E. coli*, DHBA (2000 µg/mL) combined with cefepime (512 µg/mL) or ciprofloxacin (32 µg/mL) reduced viable counts to below the detectable limit of plating under the conditions tested (Fig. 3b–c). A similar pattern was observed for the DHBA plus cefepime combination in *P. aeruginosa*, where the combination reduced bacterial recovery relative to either agent alone (Fig. 3d). Collectively, these CFU based assays confirm that DHBA can convert potentiation observed by absorbance into bactericidal outcomes with selected antibiotic partners, supporting cefepime based combinations as a particularly robust pairing for subsequent mechanistic and translational analyses.

**Fig. 3.**
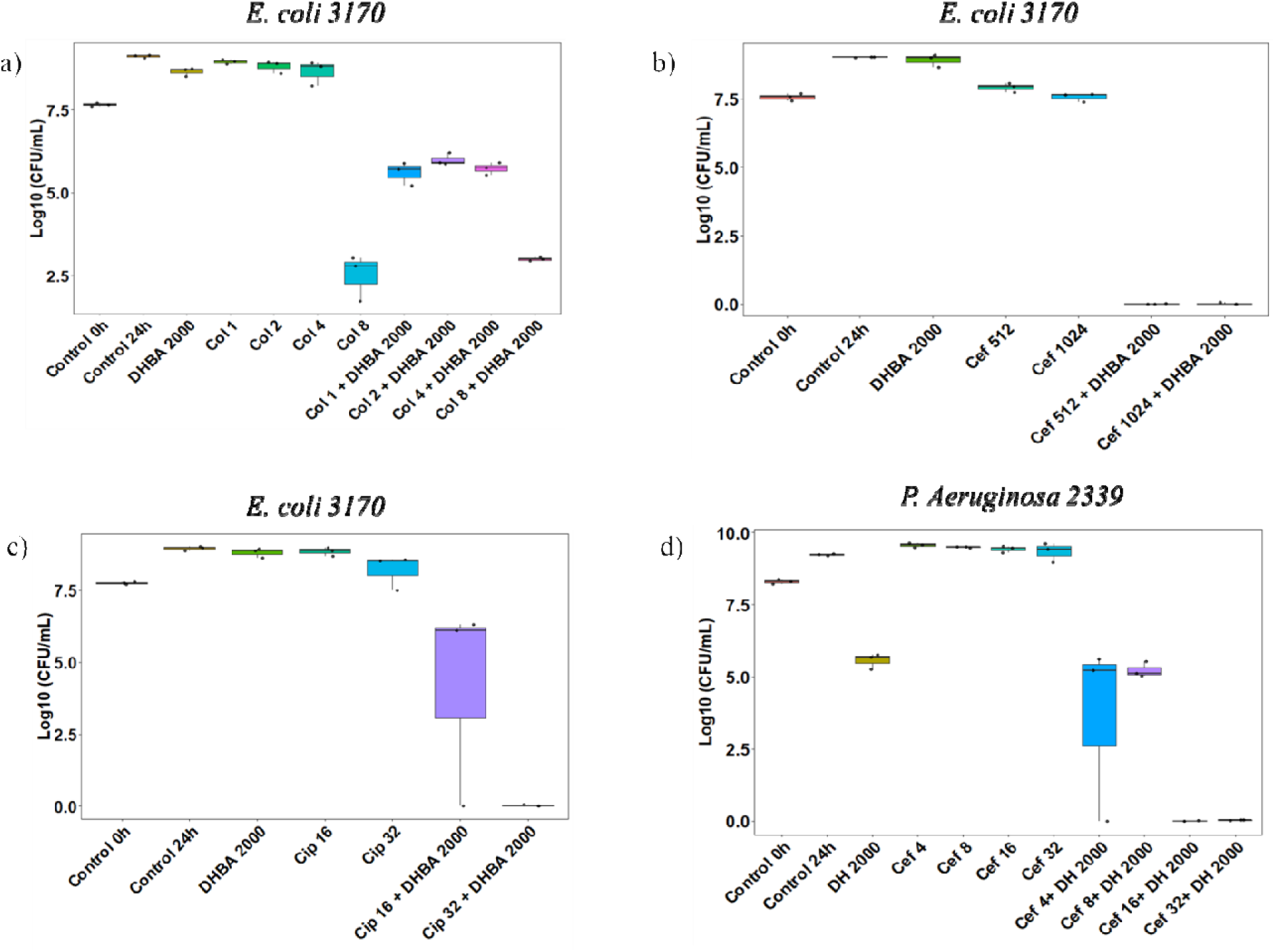
CFU enumeration confirms bactericidal activity of DHBA antibiotic combinations. Viable counts were quantified by colony forming unit (CFU) enumeration after 18 to 24 h incubation at 37 °C following exposure to DHBA, antibiotics, or their combinations, a described in Methods. (a) mcr 1 *E. coli* BAA 3170 treated with DHBA (2000 µg/mL), colistin (1 to 8 µg/mL), or the DHBA plus colistin combinations. (b) mcr 1 *E. coli* BAA 3170 treated with DHBA (2000 µg/mL), cefepime (512 µg/mL), or the DHBA plus cefepime combination. (c) mcr 1 *E. coli* BAA 3170 treated with DHBA (2000 µg/mL), ciprofloxacin (32 µg/mL), or the DHBA plus ciprofloxacin combination. (d) XDR *P. aeruginosa* 2339 treated with DHBA (2000 µg/mL), cefepime (16 µg/mL), or the DHBA plus cefepime combination. Bars represent mean log10 CFU/mL and error bars indicate standard deviation from three or more independent biological replicates. Where no colonies were recovered, values were recorded as below the limit of detection of the plating assay.

### DHBA increases outer membrane permeability and disrupts membrane integrity in E. coli

Because phenolic acids have been reported to perturb Gram negative envelopes, we tested whether DHBA alters outer membrane permeability and membrane integrity in a colistin resistant model strain, *E. coli* BAA 3170 (mcr 1). Outer membrane permeability was quantified using the 1-N phenylnaphthylamine (NPN) uptake assay. Colistin exposure increased NPN fluorescence relative to untreated cells, consistent with its established membrane active mechanism (Fig. 4a). DHBA also increased NPN fluorescence, indicating enhanced outer membrane permeability under DHBA exposure (Fig. 4a). Across the DHBA concentration series, NPN signal increased in a concentration dependent manner, supporting a direct relationship between DHBA exposure and outer membrane permeabilization (Fig. 4b). We next assessed loss of membrane integrity using propidium iodide (PI) uptake, with 70% ethanol as a positive control. Under the conditions tested, colistin produced little change in PI fluorescence relative to untreated controls, whereas DHBA increased PI uptake in a concentration dependent manner (Fig. 4c–d). These data indicate that DHBA not only increases outer membrane permeability but also compromises membrane integrity in *E. coli*. The shared NPN phenotype suggests a convergent envelope associated effect with colistin, while the distinct PI response is consistent with differences in downstream cellular damage pathways, motivating subsequent genetic susceptibility mapping to resolve DHBA specific determinants of response.

**Fig. 4.**
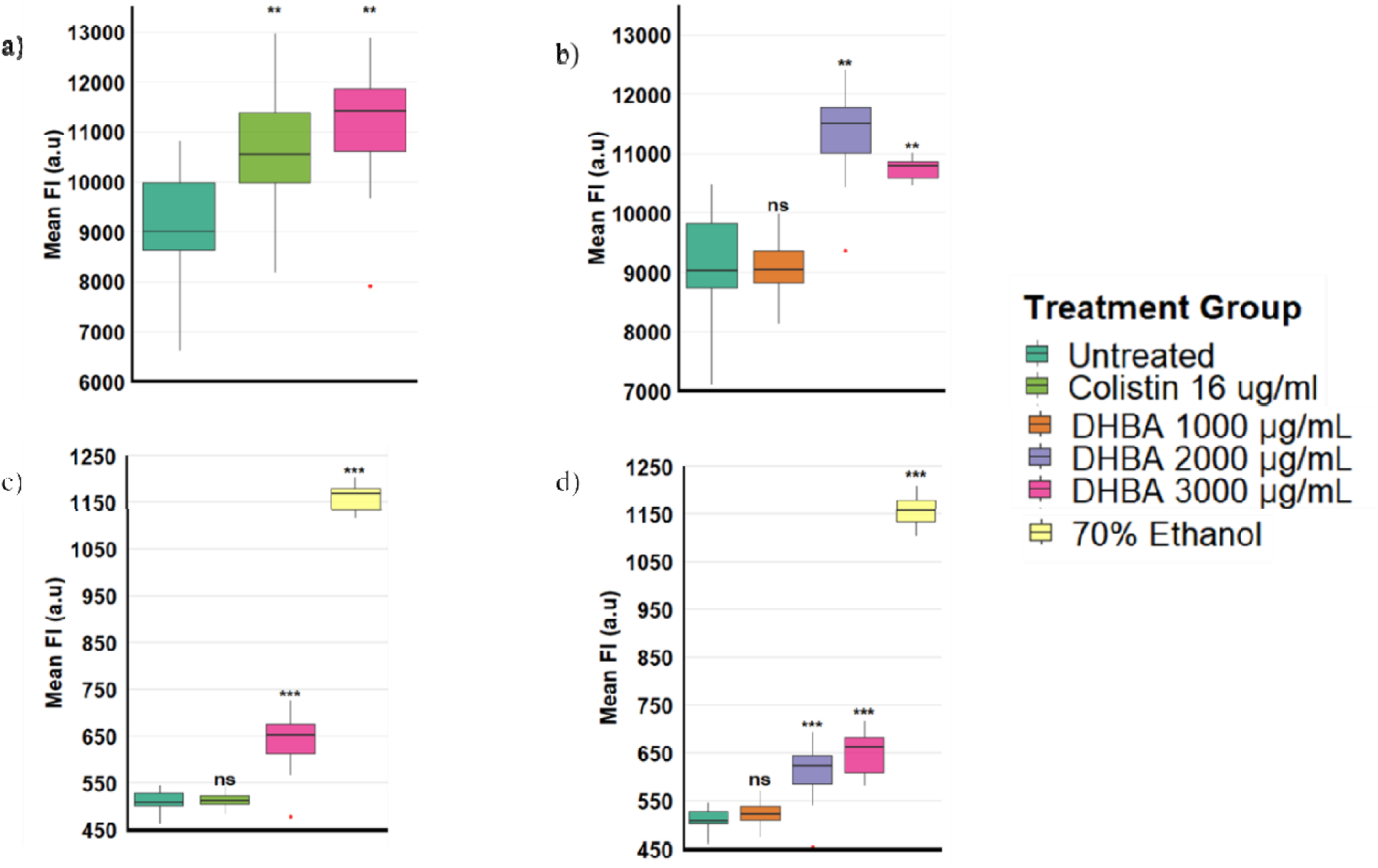
DHBA increases outer membrane permeability and disrupts membrane integrity in E. coli BAA 3170. Outer membrane permeability was assessed by 1-N phenylnaphthylamine (NPN) uptake, and membrane integrity was assessed by propidium iodide (PI) uptake, as described in Methods. (a) NPN fluorescence in *E. coli* BAA 3170 after exposure to DHBA (3000 µg/mL) or colistin (16 µg/mL) compared with untreated control. (b) NPN fluorescence after DHBA exposure at 1000, 2000, and 3000 µg/mL. (c) PI fluorescence in *E. coli* BAA 3170 after exposure to DHBA (3000 µg/mL) or colistin compared with untreated control; 70% ethanol served as a positive control. (d) PI fluorescence after DHBA exposure at 1000, 2000, and 3000 µg/mL; 70% ethanol served as a positive control. Data are presented as mean fluorescence intensity with error bars indicating standard deviation from eight independent replicates. Statistical significance relative to untreated control was evaluated using an unpaired Student’s t test (*P < 0.05, **P < 0.01).

**Fig. 5.**
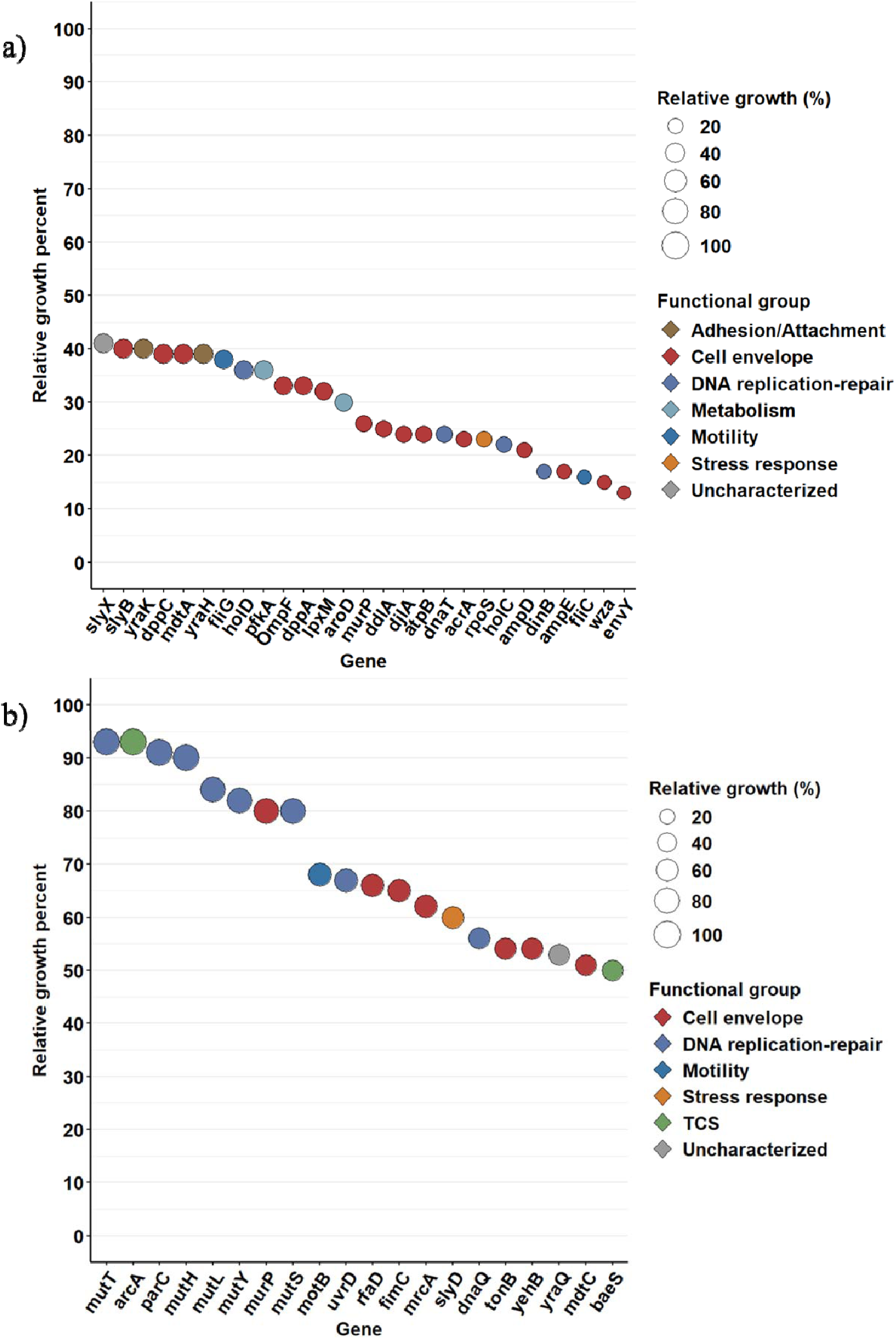
Genetic susceptibility mapping distinguishes DHBA and colistin response architectures in E. coli. Bubble plots summarizing relative growth of single gene knockout mutants screened under DHBA or colistin exposure, with mutants grouped by functional category. Bubble size denotes relative growth percentage compared with wild type *E. coli* BW25113 under the same exposure condition, as calculated from baseline and endpoint absorbance measurements (Methods). (a) Mutants showing increased relative growth under DHBA exposure (1750 µg/mL), including envelope associated determinants (for example slyB, lpxM, ompF, acrA, ampD, wza), DNA replication and repair determinants (holC, holD, dnaT, dinB), metabolic determinants (aroC, aroD, pfkA), and stress response determinants (rpoS). (b) Mutants showing increased relative growth under colistin exposure (1.5 µg/mL), including DNA replication and repair determinants (for example dnaQ, mutT, uvrD, mutH), envelope associated determinants (rfaD, tonB, mdtC, mrcA), and motility or fimbrial determinants (motB, fimC). murP was the only determinant shared between the DHBA and colistin resistant sets. Functional categories are color coded. Values represent mean relative growth in comparison to mean relative growth of WT; from four or more independent biological replicates.

### Genetic susceptibility mapping distinguishes DHBA response pathways from colistin response pathways in E. coli

DHBA and colistin differ in chemical structure and net charge, and therefore are expected to engage the Gram-negative envelope through distinct physicochemical routes. To dissect these mechanistic differences at the whole cell level, we performed a large scale genetic susceptibility screen using a curated subset of 316 single gene knockouts from the Keio collection (Supplementary Table 2). Mutants were screened under drug exposure conditions benchmarked to wild type *E. coli* BW25113, using DHBA and colistin concentrations chosen to exert substantial growth inhibition in the wild type. Mutants that showed higher relative growth than the wild type were classified as resistant in this screen and were interpreted as genetic determinants of susceptibility. Across the 316 mutant subset, DHBA resistance determinants were enriched for envelope associated functions and we identified 27 DHBA resistant mutants, and approximately half mapped to the cell envelope category (Fig:5a). Representative determinants included acrA, slyB, lpxM, dppA, dppC, mdtA, murP, ddlA, and ompF, which span lipoprotein stability, lipid A biosynthesis, efflux, peptide transport, and porin regulation. Additional envelope linked determinants included ampD, ampE, wza, and envY, consistent with a broad envelope centered response architecture. Beyond the envelope, DHBA resistance determinants also included genes involved in DNA replication and repair (holC, holD, dnaT, dinB), stress adaptation (rpoS), flagellar motility (fliG), and central metabolism (aroC, aroD, pfkA). One uncharacterized locus, slyX, also showed a marked resistant phenotype, suggesting an additional, unresolved tolerance mechanism.

In contrast, colistin resistance determinants showed a different functional distribution (Fig:5b). We identified 20 colistin resistant mutants, with a substantial fraction mapping to DNA replication and repair, including dnaQ, mutL, mutY, uvrD, mutT, mutH, and related pathways. The anaerobic regulator arcA also showed a strong resistant phenotype, consistent with links between metabolic state and colistin response [48]. Envelope related determinants were present but less dominant, including rfaD, tonB, mdtC, mrcA, yehB, fimC, and motB, which span lipopolysaccharide biosynthesis, iron transport, efflux, peptidoglycan synthesis, and motility. Additional determinants included slyD and baeS, which are linked to stress responses and envelope regulation. Notably, murP was the only gene shared between the DHBA and colistin resistant sets, suggesting a limited overlap in the genetic architecture of susceptibility despite convergent outer membrane phenotypes. We next validated selected resistant mutants using kinetic growth measurements under the same exposure conditions used in the screen. Under DHBA exposure, pfkA showed accelerated outgrowth beginning at approximately 6 h, and ompF showed progressive recovery during later time points, consistent with roles for glycolysis and porin mediated transport in shaping DHBA susceptibility (Supplementary Fig. 1a–c). Under colistin exposure, mutH and parC showed early and sustained recovery relative to the wild type, consistent with a contribution of DNA maintenance functions to colistin associated growth suppression (Supplementary Fig. 1d–f). These growth dynamics corroborate the screen and support a model in which DHBA and colistin both engage envelope associated susceptibility but diverge in downstream cellular systems that shape tolerance and recovery.

Because motility can influence adaptation to envelope stress, we further examined whether DHBA affects swimming behavior. In soft agar, DHBA reduced the motility zone diameter of the wild type strain in a concentration dependent manner, whereas the fliG mutant showed minimal change across conditions (Supplementary Fig. 2a–c). These results support flagellar function as a DHBA responsive cellular module and are consistent with the genetic susceptibility mapping that identified motility associated determinants.

### DHBA produces a more conserved community shift than colistin in an MBRA gut microbiome model

Because antibiotic exposure can destabilize resident gut microbial communities and promote outgrowth of opportunistic taxa, we next examined whether DHBA exerts collateral disruption in a complex community context. We used a Mini Bioreactor Array (MBRA) system to cultivate a stable human fecal community under continuous flow conditions. After a 7-day stabilization period, communities were treated daily for 7 days with DHBA (3000 µg/mL), colistin (6.25 µg/mL), or a DHBA plus colistin combination (2000 µg/mL DHBA with 0.4 µg/mL colistin). Endpoint samples from treated and untreated control reactors were profiled by 16S rRNA gene sequencing.

Genus level profiling revealed pronounced treatment associated restructuring across groups (Fig. 6a). Control communities retained a diverse genus distribution, with multiple taxa contributing to the overall community structure. Several genera, including *Bacteroides, Mogibacterium*, and *Anaerostipes*, were detected across conditions, although their relative abundances shifted with treatment. Colistin exposure produced a marked deviation from the control profile, including loss or strong depletion of several taxa and enrichment of opportunistic genera. In particular, colistin treated communities showed depletion of *Anaerococcus* and a reduction in *Bacteroides*, accompanied by enrichment of *Escherichia-Shigella*. Several genera absent or low in controls increased under colistin exposure, including *Dorea, Lachnoclostridum, Lactobacillus*, and *Clostridium sensu stricto*.

**Fig. 6.**
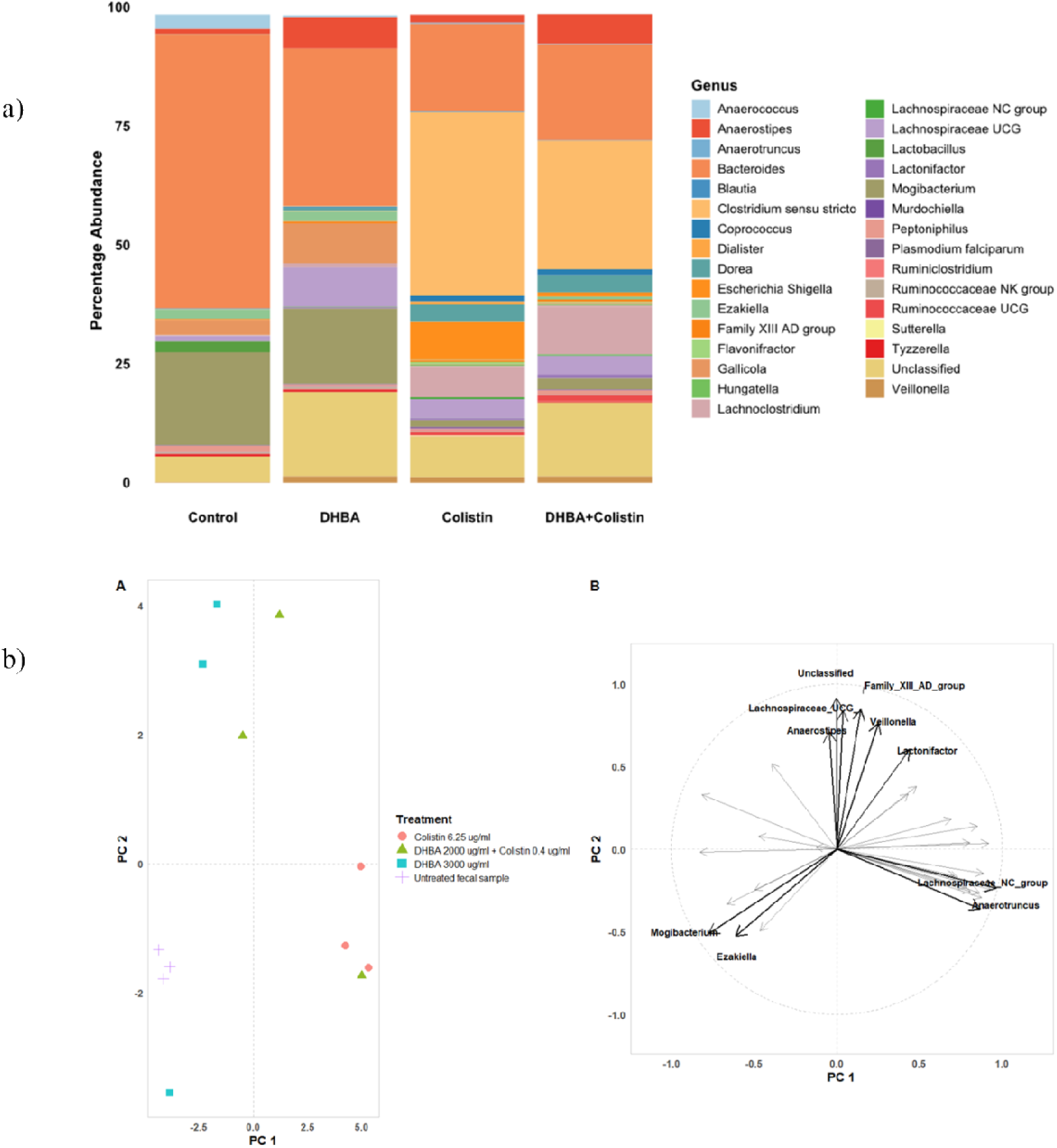
Community level effects of DHBA and colistin in an MBRA gut microbiome model. Human fecal communities were cultivated in a Mini Bioreactor Array system, stabilized for 7 days, and then treated daily for 7 days with DHBA (3000 µg/mL), colistin (6.25 µg/mL), or the DHBA plus colistin combination (DHBA 2000 µg/mL with colistin 0.4 µg/mL), as described in Methods. Endpoint samples were profiled by 16S rRNA gene sequencing. (a) Genus level relative abundance profiles for each treatment group. (b) Principal coordinates analysis (PCoA) of community structure based on Bray Curtis dissimilarity, showing separation of samples by treatment condition. Arrows indicate genera contributing to variation along the principal coordinate axes in the biplot.

In contrast, DHBA treatment produced a more conserved shift relative to colistin. Multiple genera present in the control, including *Anaerococcus, Flavonifractor, Peptoniphilus,* and *Gallicola*, were retained under DHBA exposure, although relative abundance changes were observed. DHBA treated communities showed increased relative abundance of several taxa often associated with anaerobic fermentation, including *Blautia, Anaerostipes,* and *Lachnospiraceae*, alongside reduced abundance of *Anaerococcus* and *Bacteroides*. In the DHBA plus colistin condition, community profiles more closely resembled the colistin treated group, indicating that colistin exposure remained a dominant driver even at the lower concentration tested. However, the combination showed reduced enrichment of opportunistic genera relative to colistin alone, including attenuation of *Clostridium sensu stricto, Escherichia* and *Shigella*. Across all groups, a substantial fraction of reads remained assigned to unclassified taxa, consistent with incomplete representation of community members in reference databases. Ordination analysis demonstrated distinct clustering by treatment (Fig. 6b). Control reactors clustered tightly, whereas treated communities formed separate clusters, indicating consistent, condition specific shifts in community structure. The DHBA and colistin groups separated from controls, and the combination group formed a distinct cluster associated with altered abundance of genera including *Peptoniphilus, Tyzzerella, Veillonella,* and *Ruminococcaceae*. Collectively, these data indicate that colistin exposure strongly remodels community composition and is associated with enrichment of opportunistic taxa, whereas DHBA produces a more conserved community shift and can partially attenuate specific colistin associated enrichments in the combination condition.

### DHBA antibiotic combinations improve efficacy in infection relevant in vivo and ex vivo models

To determine whether DHBA mediated potentiation observed in vitro translates to infection contexts, we evaluated DHBA antibiotic combinations in complementary infection relevant models (Fig. 7). DHBA showed no signs of toxicity when tested in uninfected worms. In a *G. mellonella* survival model infected with *E. coli* BAA 3170, treatment at 1 h post infection with DHBA plus colistin (500 mg/kg DHBA with 2.5 mg/kg colistin; n = 17 larvae) improved survival over the 5-day monitoring period relative to colistin alone (n = 20) and untreated controls (n = 17) (Fig. 7a; P = 0.0225 for combination versus untreated).

**Fig. 7.**
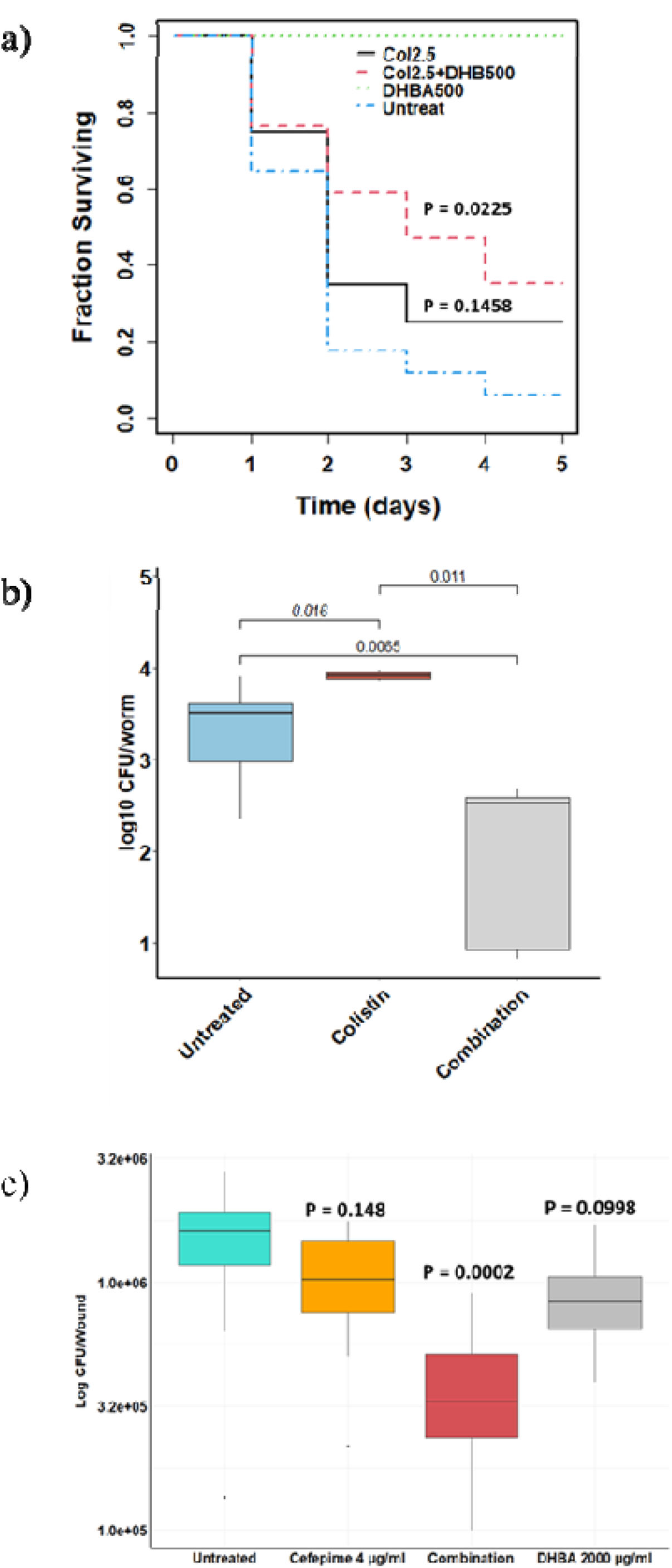
DHBA antibiotic combinations improve efficacy in infection relevant in vivo and ex vivo models. (a) Kaplan Meier survival of G. mellonella larvae with DHBA (no infection), infected with *E. coli* BAA 3170 and treated 1 h post infection with colistin (2.5 mg/kg; n = 20), DHBA plus colistin (DHBA 500 mg/kg with colistin 2.5 mg/kg; n = 17), or left untreated (n = 17). Survival differences were evaluated by log rank test; DHBA plus colistin improved survival relative to colistin alone. (b) Intestinal bacterial burden in Caenorhabditis elegans after exposure to DHBA plus colistin (DHBA 2000 µg/mL with colistin 1 µg/mL), quantified as CFU per worm. (c) Bacterial burden in an ex vivo porcine skin burn wound infection model infected with XDR *P. aeruginosa* 2339 after treatment with DHBA (2000 µg/mL), cefepime (4 µg/mL), or the DHBA plus cefepime combination (n = 10 wounds per condition). Wound burden is reported as log CFU per wound.

We next assessed intestinal colonization burden in *C. elegans*. Exposure to DHBA plus colistin (2000 µg/mL DHBA with 1 µg/mL colistin) reduced internal *E. coli* burden at the endpoint relative to untreated controls (Fig. 7b), indicating improved pathogen clearance under host associated conditions. Finally, we tested DHBA potentiation in a clinically relevant local infection setting using an ex vivo porcine skin burn wound model infected with XDR *P. aeruginosa* 2339. Treatment with DHBA plus cefepime (2000 µg/mL DHBA with 4 µg/mL cefepime) reduced wound associated bacterial burden relative to untreated controls and to single agent conditions (Fig. 7c).

## Discussion

### Systems level mapping and mechanistic model

Antibiotic action in Gram-negative bacteria is increasingly recognized as an emergent property of envelope access, stress and repair networks, and metabolic state, rather than a single primary target alone(33–35). In this study, we apply a systems level mapping framework to a microbiota derived phenolic acid potentiator by integrating potentiation assays, CFU enumeration, membrane permeability and integrity measurements, and genetic susceptibility profiling across major cellular modules. We used colistin as a comparator to separate convergent envelope associated phenotypes from divergent downstream susceptibility architectures. This framework provides a mechanistic basis for interpreting class selective potentiation and for prioritizing combinations for infection relevant settings. Membrane phenotyping provided direct evidence that DHBA alters envelope properties in a resistant Gram-negative background. DHBA increased NPN uptake, indicating enhanced outer membrane permeability. In contrast to colistin under the conditions tested, DHBA also increased propidium iodide uptake in a concentration dependent manner, consistent with disruption of membrane integrity. These phenotypes support a model in which DHBA induces an envelope permissive state that can increase access of partner antibiotics to their sites of action. This interpretation is consistent with the broader literature on phenolic acids, where membrane associated perturbations are frequently proposed, but are often not mechanistically resolved(19, 36, 37).

Our genetic susceptibility mapping using single gene mutants adds an additional layer of mechanistic resolution by identifying cellular systems that shape DHBA responsiveness. Resistant determinants were enriched for envelope associated functions, including porin regulation, efflux, lipoprotein stability, and lipid A associated processes, which is consistent with the permeability phenotypes. However, the mapping also identified determinants outside the envelope, including central metabolism and information processing functions. These findings argue against a single locus mechanism and instead support a multi module susceptibility architecture in which envelope perturbation is a primary feature, while metabolic state and stress responses shape tolerance and recovery(38–41). The growth kinetics validation aligns with this concept, as mutants in glycolysis and porin related loci showed distinct recovery dynamics under DHBA exposure.

Colistin provides a useful comparator because it is a canonical membrane active antibiotic and remains clinically relevant in resistant Gram-negative infections(42–44). DHBA and colistin shared a convergent outer membrane phenotype by NPN uptake, but they diverged in propidium iodide uptake under the conditions tested and, more importantly, in genetic susceptibility architecture. DHBA resistance determinants were dominated by envelope and transport systems with additional metabolic and replication linked factors, whereas colistin resistant determinants were comparatively enriched for DNA maintenance and repair functions alongside a smaller envelope component. The limited overlap between the DHBA and colistin determinant sets, with murP as the primary shared locus, suggests that the two conditions impose different cellular pressures despite a shared envelope associated entry phenotype. This distinction matters for combination design, because potentiators that act through a susceptibility architecture distinct from the partner antibiotic may provide complementary leverage and may reduce direct mechanistic redundancy. More broadly, these results support the concept that both potentiators and antibiotics can engage secondary cellular systems beyond canonical primary mechanisms, and that mapping these determinants can reveal actionable constraints for therapy design.

### Implications for combination design and translation

A central functional outcome of our study is that DHBA restores susceptibility to selected antibiotic classes in resistant Gram-negative strains, including mcr 1 positive *E. coli* and XDR clinical isolates. The sixteen-fold reduction in colistin MIC in mcr 1 positive *E. coli* is notable because it represents a shift in susceptibility for a last resort antibiotic under conditions where resistance otherwise compromises therapy. DHBA also restored measurable activity of cefepime and ciprofloxacin in the same *E. coli* background, and potentiation extended to XDR *P. aeruginosa* and *K. pneumoniae* in antibiotic specific combinations. Importantly, potentiation was not universal. DHBA did not improve kanamycin activity, which provides an internal specificity control and helps constrain mechanistic interpretation.

The kanamycin result fits the systems level framework described above. Aminoglycosides require entry through the envelope and subsequent energy dependent transport across the inner membrane, and activity is sensitive to membrane potential and metabolic state. The absence of meaningful potentiation for kanamycin indicates that DHBA does not uniformly enhance uptake or intracellular access for all antibiotic classes. Instead, potentiation appears stronger for antibiotics whose activity can benefit from altered envelope permeability or periplasmic access, including polymyxins and beta lactams, and for ciprofloxacin in the tested resistant background. This class selectivity is consistent with a defined susceptibility reshaping mechanism rather than broad growth suppression, and it provides an interpretable rule for prioritizing partner antibiotics(45, 46). In practical terms, it suggests that envelope access and downstream recovery pathways should be treated as design constraints when selecting combinations, rather than assuming that any antibiotic will benefit from a potentiator that perturbs the envelope. CFU based assays further support the functional relevance of DHBA potentiation. In *E. coli*, DHBA increased colistin associated killing at lower colistin exposures and enabled clearance to below the detectable limit of plating when paired with cefepime or ciprofloxacin under the tested conditions. The DHBA cefepime combination also reduced recovery of XDR *P. aeruginosa*.

Collateral disruption of commensal microbiota is another dimension of translational relevance, particularly when potentiators are intended to extend antibiotic use(47–49). In an MBRA model of a human fecal community, colistin exposure produced marked restructuring of genus level composition and enrichment of opportunistic taxa, including *Escherichia-Shigella*, alongside depletion of several genera detected in control communities. DHBA exposure also shifted community composition, but the resulting profiles were more conserved relative to colistin, with retention of several genera observed in controls, including *Flavonifractor,* and enrichment of taxa often associated with anaerobic fermentation, such as *Blautia and Anaerostipes*. In the DHBA plus colistin condition, community profiles resembled the colistin group, indicating that colistin remained a dominant driver even at the lower concentration used. However, DHBA attenuated selected colistin associated enrichments in opportunistic taxa. These observations suggest that DHBA may offer a narrower community level perturbation profile than colistin under the tested conditions, and they motivate future work that links mechanistic susceptibility determinants to community level outcomes.

Potentiation observed *in vitro* can fail to translate under host associated conditions due to matrix effects, physiological constraints, or altered bacterial physiology. We therefore evaluated DHBA antibiotic combinations in infection relevant models spanning distinct contexts. In *G. mellonella* infected with *E. coli* BAA 3170, DHBA plus colistin improved survival relative to colistin alone, supporting a functional impact in a whole organism setting. In *C. elegans*, DHBA plus colistin reduced internal bacterial burden in an intestinal colonization context. Finally, in an *ex vivo* porcine skin burn wound model infected with XDR *P. aeruginosa*, DHBA plus cefepime reduced wound associated bacterial burden relative to controls and single agent conditions. Together, these models provide convergent evidence that DHBA potentiation is retained under infection like conditions and that DHBA cefepime combinations are especially relevant in local infection contexts where high local concentrations are practical.

### Limitations and next steps

Out study has several limitations. First, DHBA activity and potentiation were observed at high concentrations, and translation will likely depend on formulation and route of delivery. The porcine skin model supports feasibility for topical or local applications, but additional work is required to evaluate tissue compatibility and wound healing effects under repeated exposure. Second, genetic susceptibility mapping identifies determinants that shape whole cell response, but does not establish direct molecular binding targets. Complementation assays and orthogonal biochemical studies will be needed to convert susceptibility determinants into validated target mechanisms and to clarify whether the strongest determinants reflect entry, efflux, stress adaptation, or other coupled processes. Third, although the multi module susceptibility architecture suggests a complex pressure landscape, this study does not measure resistance emergence rates under DHBA antibiotic combinations. Serial passaging and frequency of resistance assays will be required to test whether combinations increase the genetic barrier to resistance in specific strain backgrounds. Finally, MBRA results were generated in a defined in vitro community context, and broader validation across donors and dosing schedules will be needed to establish general microbiome outcomes.

In summary, DHBA represents a microbiota derived phenolic acid scaffold that can restore susceptibility to selected antibiotic classes in resistant Gram-negative pathogens. By integrating potentiation assays, CFU based killing, membrane phenotyping, and system wide genetic susceptibility mapping, this work provides a mechanistic framework to guide rational optimization of DHBA based combinations and to prioritize pairing strategies for further development, particularly in local infection contexts where high local exposure is feasible.

## Acknowledgments

This work was supported in part by the Walter R. Sitlington Endowed Chair in Infectious Diseases. This material is based upon work supported by the United States Department of Agriculture, National Institute of Food and Agriculture (USDA NIFA). We thank the CDC ARLG Antibiotic Resistance Isolate Bank for providing bacterial isolates used in this study. We thank Prabhjot Kaur Sekhon for assistance with Mini Bioreactor Array experiments. We also thank the donors who provided fecal samples for MBRA studies.

---

**Supplementary Fig. 1.**
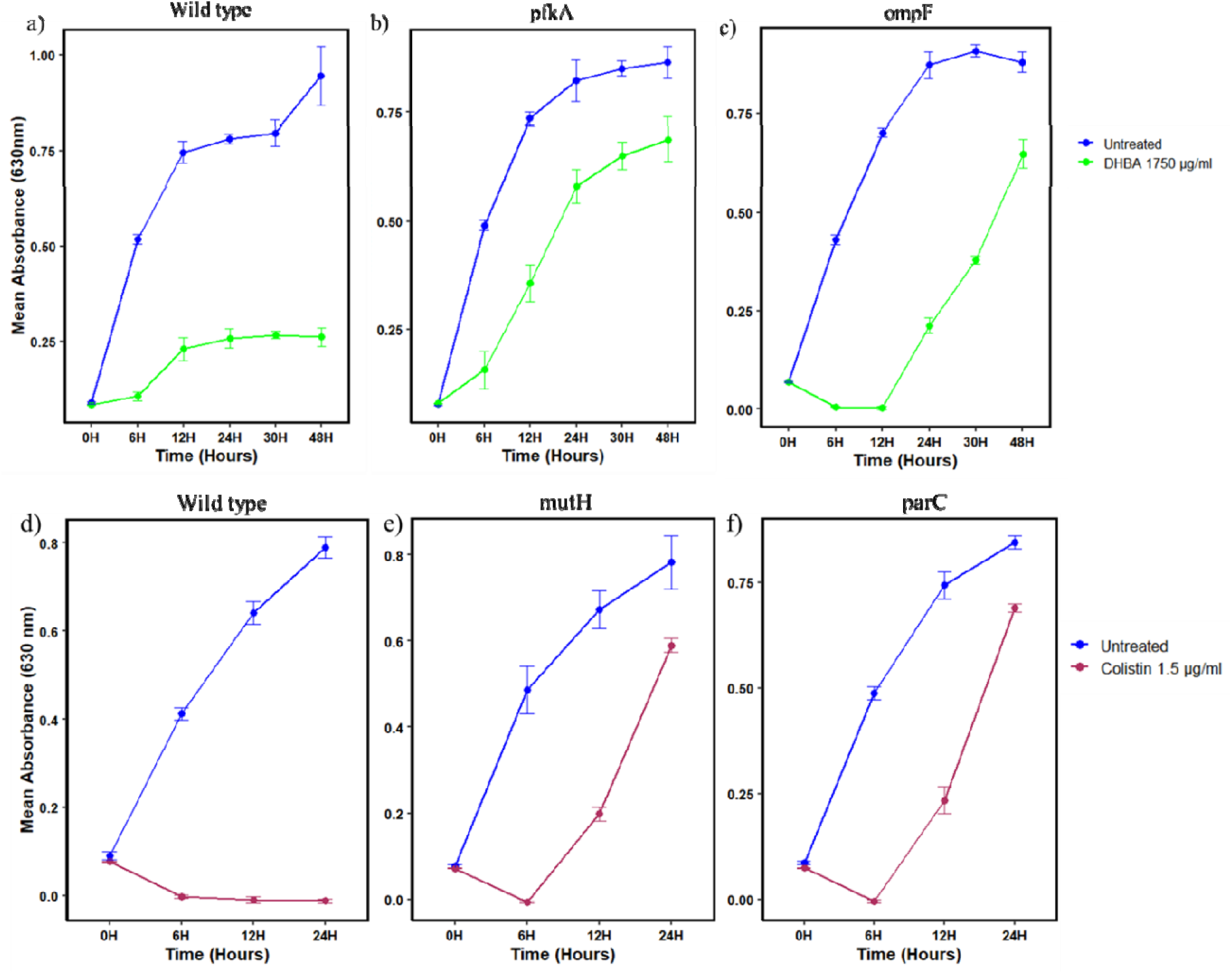
Kinetic growth validation of resistant E. coli mutants under DHBA and colistin exposure. (a–c) Growth kinetics of wild type E. *coli* BW25113 and the pfkA and ompF knockout mutants during exposure to DHBA (1750 µg/mL), showing accelerated outgrowth of pfkA and delayed recovery of ompF relative to the wild type. (d–f) Growth kinetics of wild type *E. coli* BW25113 and the mutH and parC knockout mutants during exposure to colistin (1.5 µg/mL), showing early and sustained recovery of mutH and parC relative to the wild type. Optical density (OD630) was recorded at regular intervals during incubation, as described in Methods.

**Supplementary Fig. 2.**
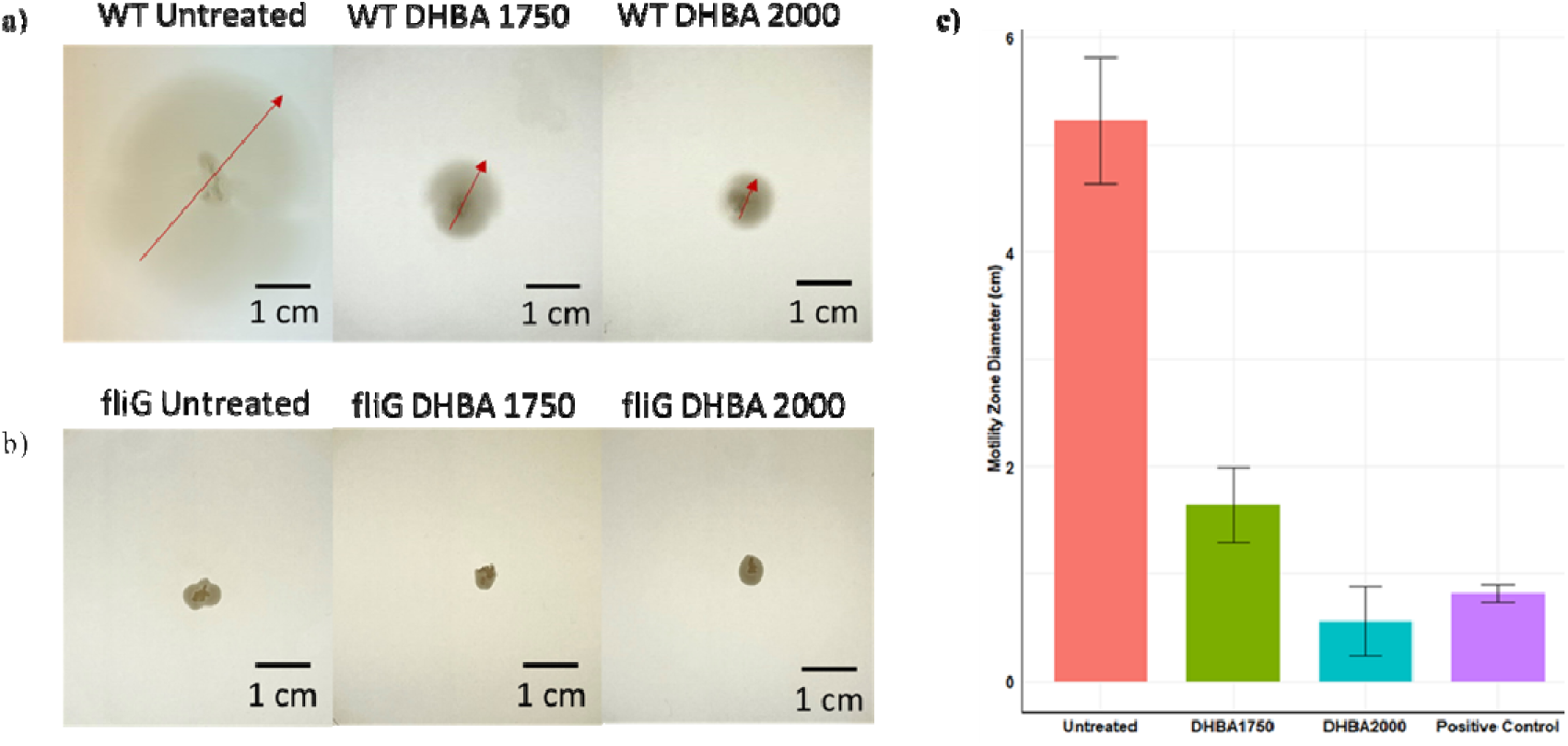
DHBA reduces swimming motility in wild type E. coli but not in a fliG mutant. (a–b) Representative LB soft agar (0.3% agar) swimming plates showing migration zones of wild type *E. coli* and the fliG knockout mutant under untreated conditions and after DHBA exposure. Red arrows indicate the edge of the motility zone. Scale bar = 1 cm. (c) Quantification of motility zone diameter across conditions, including untreated wild type, wild type treated with DHBA (1750 µg/mL or 2000 µg/mL), and the fliG mutant. Wild type motility decreased with DHBA exposure, whereas the fliG mutant showed minimal change across conditions. Values represent mean diameter with error bars indicating standard deviation from five independent biological replicates.

